# Single cell transcriptional profiling reveals helper, effector, and regulatory MAIT cell populations enriched during homeostasis and activation

**DOI:** 10.1101/2020.10.22.351262

**Authors:** Charles Kyriakos Vorkas, Chirag Krishna, Kelin Li, Jeffrey Aubé, Daniel W. Fitzgerald, Linas Mazutis, Christina S. Leslie, Michael S. Glickman

## Abstract

Mucosal-associated invariant T (MAIT) cells are innate-like lymphocytes that recognize microbial vitamin B metabolites and have emerging roles in infectious disease, autoimmunity, and cancer. Although MAIT cells are identified by a semi-invariant T cell receptor, their phenotypic and functional heterogeneity is not well understood. Here we present an integrated single cell transcriptomic analysis of over 76,000 human MAIT cells during acute and chronic antigen-specific activation with the MR1 ligand 5-OP-RU and non-specific TCR stimulation. We show that MAIT cells span a broad range of homeostatic, effector, helper, tissue-infiltrating, regulatory, and exhausted phenotypes, with distinct gene expression programs associated with CD4^+^ or CD8^+^ co-expression. During acute activation, MAIT cells rapidly adopt a cytotoxic phenotype characterized by high expression of *GZMB*, *IFNG* and *TNF*. In contrast, chronic stimulation induces heterogeneous states defined by proliferation, cytotoxicity, immune modulation, and exhaustion. These scRNAseq-defined MAIT cell subtypes were also detected in individuals recently exposed to *Mycobacterium tuberculosis* infection, confirming their presence during human infection. Our study provides the first comprehensive atlas of human MAIT cells in activation conditions and defines substantial functional heterogeneity, suggesting complex roles in health and disease.

## Introduction

Mucosal-associated invariant T (MAIT) cells are a subset of re-circulating innate-like lymphocytes enriched in human liver, lung and gut with highly variable abundance in peripheral blood in healthy donors (<1%-18% in humans)(Godfrey et al., 2019; Li et al., 2018; Vorkas et al., 2018). MAIT cells express an evolutionarily conserved T cell receptor (TCR; TRAV1-2 in humans) that can recognize microbially-derived, non-peptide small molecule metabolites of the vitamin B pathway(Harriff et al., 2018; Treiner et al., 2003) presented by the oligomorphic MHC Ib-related molecule, MR1(Seshadri et al., 2017; Treiner et al., 2003). In contrast to conventional T cells that exit the thymus in a “naive” state and require peptide-specific antigen priming over days to weeks to proliferate, most mature MAIT cells enter the circulation as effector cells and are activated within hours of antigen recognition(Legoux et al., 2019). This innate-like property confers their capacity to be among the first responders to bacterial and fungal pathogens(Dias et al., 2017; Jahreis et al., 2018; Shaler et al., 2017; Vorkas et al., 2018) as well as to viral infections through TCR-independent cytokine stimulation (Lamichhane et al., 2019; Leng et al., 2019; van Wilgenburg et al., 2016).

Despite a growing body of literature implicating MAIT cells in both homeostasis and disease(Godfrey et al., 2019), their specialized functional states in humans are poorly characterized relative to conventional T cells. For example, whereas conventional T cells are broadly divided into two lineages defined by CD4 or CD8 co-receptor expression and stable functional phenotypes, whether these same lineages can be defined in MAIT cells remains unclear. CD4^+^ MAIT cells are reported to express increased CD25 and IL2 along with decreased cytotoxic markers relative to CD8^+^ MAIT cells (Gherardin et al., 2018b; Kurioka et al., 2017; Lamichhane et al., 2019; Vorkas et al., 2018). While CD8^+^ and CD4^−^CD8^−^ (double-negative; DN) MAIT cells have similar functional phenotypes(Kurioka et al., 2017), the DN subset is reported to have depressed cytotoxic function with increased propensity to activation-induced apoptosis(Dias et al., 2018). Moreover, while prior studies have suggested that MAIT cell functional states may depend on the quality and duration of activating stimuli(Kelly et al., 2019; Lamichhane et al., 2019), the early and late effects of TCR-stimulation on MAIT cell state remain largely unknown.

Here, we present a single cell transcriptomic atlas of human MAIT cells, comprising more than 76,000 cells from peripheral blood under acute and chronic activation with 5-OP-RU or αCD3/CD28 in the presence of IL2 or IL2/TGFβ. Our study elucidates distinct transcriptional programs of CD4^+^ and CD8^+^ MAIT cells and identifies novel MAIT cell functional clusters that we validate in human subjects recently exposed to *Mycobacterium tuberculosis*. Our results demonstrate broad functional heterogeneity of MAIT cells and suggest that these innate T cells can participate in complex immune responses in addition to their role in direct killing of pathogens.

## Results

### CD4 or CD8 expression on MAIT cells is associated with distinct transcriptional states

To simulate acute and chronic MAIT cell activation under antigen-specific and non-specific T cell receptor stimulation, we employed an in vitro activation assay using human PBMCs incubated with MR1 ligand (5-OP-RU) or αCD3/CD28 beads for 15 hours (D1; acute) or 7 days (D7; chronic) compared to unstimulated control cells. A subset of conditions at D7 also included TGFβ to further assess potential regulatory phenotypes(Bhattacharyya et al., 2018; Rouxel et al., 2017). Cells were harvested at each time point for flow cytometric assays and flow-cytometry assisted cell sorting (FACS) using MR1 tetramers (Fig. 1a, Supplementary Fig. 1, and Supplementary Table 1). To define the immunologic landscape of MAIT cell states after acute and chronic activation, we performed droplet microfluidics single cell RNA sequencing of these sorted MAIT cells from the peripheral blood of three donors in the activation conditions defined above. 76,845 MAIT cells passed quality control and were used for downstream analysis.

**Figure 1.**
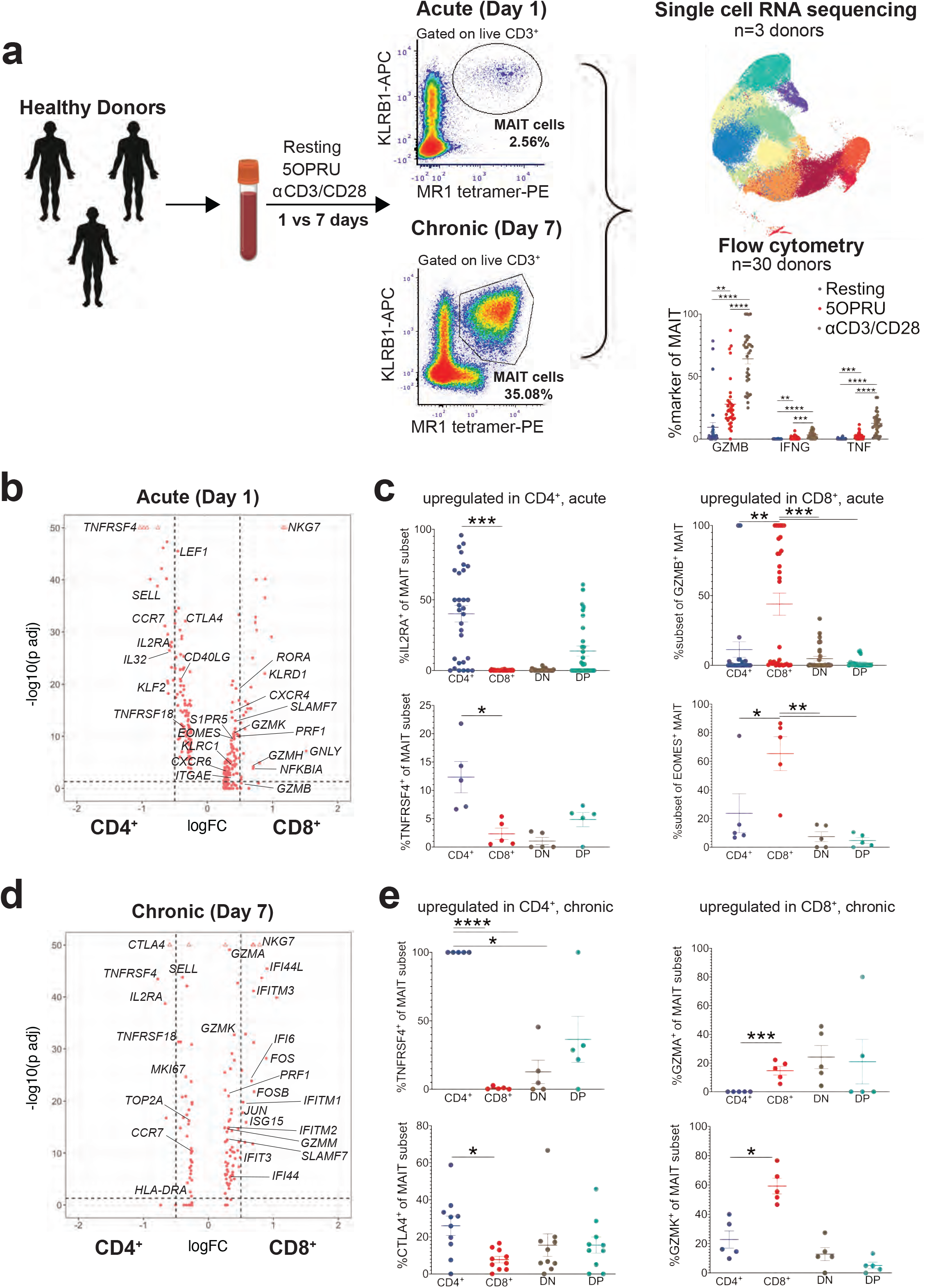
MAIT cell CD4 or CD8 expression are associated with distinct transcriptional signatures. **(a)** Experimental schematic of the MAIT cell in vitro activation assay used for single cell RNA sequencing (scRNA-seq) and flow cytometric analyses. A total of 30 healthy donor PBMCs were assayed by flow cytometry after 15 hrs (D1; acute activation) or 7 days (D7; chronic activation) of incubation under resting, 5OPRU or αCD3/CD28 conditions. A subset of three of these donors underwent flow cytometry-assisted cell sorting (FACS) for scRNA-seq under the same conditions. All conditions included IL2 and a subset of D7 conditions also included TGFβ. **(b)** Differentially expressed genes in CD8^+^ (+ log FC; log fold change) or CD4^+^ MAIT cells (−log FC) after 15 hours of incubation in the resting condition. y axis: log FDR-adjusted p value with a significance level of FDR < 0.05. **(c)** Mean% +/− SEM of CD25, OX40, GZMB, and EOMES staining in CD4^+^, CD8^+^, DN or DP MAIT cell subsets measured by flow cytometry in the same condition as (b). Data representative of 3 independent experiments. n=5-30 donors. **(d)** Differentially expressed genes in CD8^+^ or CD4^+^ MAIT cells after 7 days of incubation in the resting condition. Same axes as (b). **(e)** Mean% +/− SEM of OX40, CTLA4, GZMA, and GZMK staining in MAIT cell subsets measured by flow cytometry in the same condition as (d). Data representative of 2 independent experiments. n=5-10 donors. Flow cytometric statistical comparisons were made by unpaired t-test. *p<0.05 **p<0.005 *** p<0.0005 ****p<0.0001; GZMB: granzyme B; GZMA: granzyme A; GZMK: granzyme K; DN: double-negative (CD4^−^CD8); DP: double-positive (CD4^+^CD8^+^)

Although prior studies have demonstrated that MAIT cells can express CD4 or CD8 co-receptors(Gherardin et al., 2018b; Kurioka et al., 2017; Vorkas et al., 2018), it is unclear whether these markers define functionally distinct MAIT cell populations. We first sought to resolve this question using our single cell data by performing a differential expression analysis between CD4^+^CD8^−^ and CD4^−^CD8^+^ MAIT cells at D1 (Fig. 1b,c) or D7 (Fig. 1d, e) in the absence of TCR stimulation. At D1 (Fig. 1c) CD4^+^ MAIT cells more highly express genes associated with conventional naïve and central memory T cells including *LEF1*, *SELL*, and *CCR7*, but also markers of activation and co-stimulation such as *CTLA4*, *IL2RA*, *TNFRSF4*, and *TNFRSF18* (Fig 1b, Supplementary Table 2). CD8^+^ MAIT cells more highly express markers of cytotoxic activity such as *NKG7*, *KLRD1*, *EOMES*, *GNLY*, and *GZMB* as well as the transcription factor *RORA*.

We confirmed the differential expression of CD25, OX40, GZMB, and EOMES at the protein level within CD4^+^, CD8^+^, DN and CD4^+^CD8^+^ (double-positive; DP) MAIT cell subsets using flow cytometry (Fig. 1c). At D7, CD4^+^ MAIT cells upregulated *CTLA4*, *TNFRSF4*, *SELL*, *IL2RA* and *CCR7*, whereas CD8^+^ MAIT cells upregulated *NKG7*, *GZMA*, *GZMK*, *PRF1*, SLAMF7 as well as several genes involved in Type I interferon signaling (*IFI44L*, *IFITM3*, *IFI6*, *IFITM1*, and *ISG15*) (Fig. 1d, Supplementary Table 3). We confirmed the differential expression of OX40, CTLA4, GZMA, and GZMK in each subset at the protein level (Fig. 1e). Our analysis demonstrates that CD4^+^ MAIT cells are associated with expression of co-stimulatory receptors, IL2 signaling and memory markers, whereas CD8^+^ MAIT cells are defined by granzyme-mediated cytotoxicity and type 1 interferon signaling. These results suggest that CD4 and CD8 expression on MAIT cells may define distinct functional roles.

### Single cell transcriptional analysis defines heterogeneous MAIT cell activation states

To fully characterize the phenotypic states of tetramer positive MAIT cells, we performed dimensionality reduction by principal component analysis followed by Louvain clustering on cells from all donors, timepoints, and stimuli (Fig. 2a). Clusters were broadly partitioned by timepoint (Supplementary Fig. 2). Visualization with uniform manifold approximation and projection (UMAP) together with differential expression analysis displayed 12 clusters that could be distinguished by homeostatic, cytotoxic, Th1-associated cytokine release, proliferating, tissue-infiltrating, immune regulatory, and exhausted states (Fig. 2b-c; Supplementary Table 3). Clusters 1-3 were enriched at D1 (acute activation). Cluster 1 (C1) had high expression of *IFNG*, *GZMB*, *TNF*, *TBX21*, and notably the transcription factor *FOXP3*. Since this cluster expressed genes associated with granzymes and Th1-associated cytokines, it was termed “MAIT1 cytotoxic effector” (Koay et al., 2019), consistent with a similar cluster detected in Koay *et al.*(Koay et al., 2019) C2 demonstrated increased *CD69* and pronounced expression of heat shock proteins, *NR4A1*, and the AP-1 subunits *JUN* and *FOS* and was named “Early activated”. C3 expressed effector/central memory markers *CCR7* and *C62L*, markers of MAIT cell intrathymic development *CD4* and *LEF1*(Koay et al., 2019), as well as *RUNX3*. This cluster also upregulated type 1 interferon signaling genes, e.g. *IFIT3* and *ISG15*. Thus, C3 was named “ISG^+^ memory” (ISG: interferon-stimulated genes). Collectively these clusters define the landscape of MAIT cells during homeostasis and early activation. The rapid induction of cytotoxic effector programs at this timepoint suggest that MAIT cells may be poised to respond earlier than conventional T cells.

**Figure 2.**
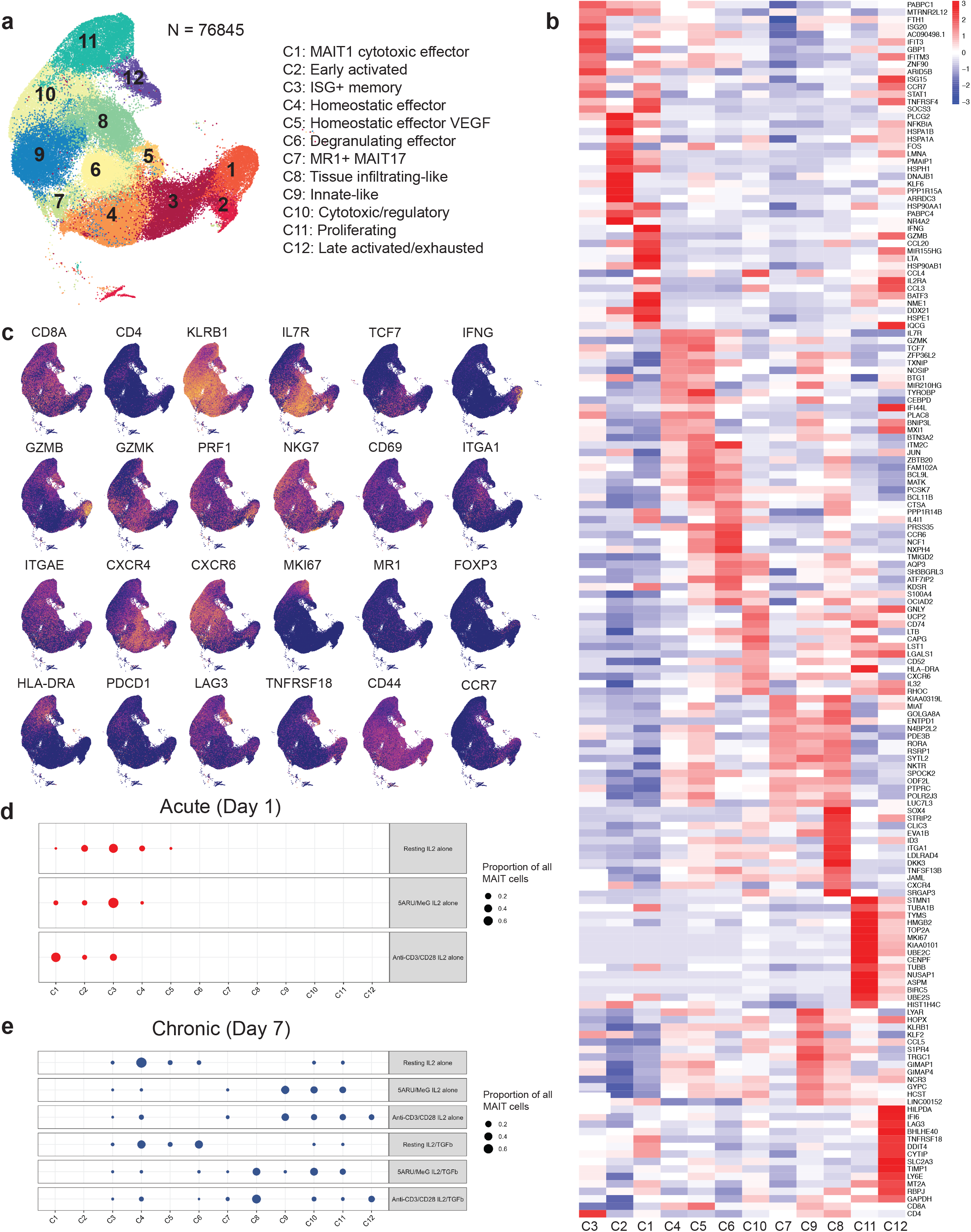
Single cell transcriptional analysis defines heterogeneous MAIT cell activation states. **(a)** Uniform manifold approximation and projection (UMAP) plot and Louvain clustering of cells from all donors, timepoints and stimuli with ascribed names based on differentially expressed gene sets in each cluster. **(b)** Heat map showing top 15 differentially expressed genes per cluster after unsupervised hierarchical clustering. **(c)** UMAP plots of selected MAIT cell markers demonstrating expression level across all clusters (blue=low expression to yellow=high expression). **(d, e)** Dot plots demonstrating relative frequency of cells per condition in all donors.

In contrast, clusters 4-12 were enriched with chronic activation and demonstrated marked phenotypic heterogeneity. C4 and C5 (“Homeostatic effectors”) displayed similar expression patterns, with high levels of *IL7R*, *GZMM*, *GZMK*, *KLRG1* and *IFI44L* and decreased expression of *CD69* and *CD25*. C4 was distinguished from C5 by expression of vascular endothelial growth factors *VEGFA/B* and was named “Homeostatic effector VEGF.” C6 (“Degranulating effector”) showed high expression of *LAMP1*, cytokines *TNF* and *TNFSF13B* (B-cell activating factor), *IFNGR1*, as well as the homing molecule *CXCR3*. C7 (“MR1^+^ MAIT17”) expressed *GZMA*, *ENTPD1*, *RORA*(Koay et al., 2019) as well as *MR1* itself, suggesting that this population may interact with other MAIT cells. C8 (“Tissue-infiltrating-like”) highly expressed *ITGA1* and *ITGAE* and was only present in activation conditions including TGFβ. C9 was termed “Innate-like” given its high expression of *HOPX*, *KLRB1*, and *NCR3* together with *GZMA* and *GZMK*. Clusters 10-12 were defined by a gradient of activation/exhaustion marker expression including *PDCD1*, *CTLA4*, *PDCD1* and *LAG3*, with C12 (“Late activated/exhausted”) resembling a terminally exhausted population. C11 (“Proliferating”) demonstrated a strong mitotic phenotype. C10 (“Cytotoxic/Regulatory”) upregulated *GZMA*, *GNLY* and *IL26* as well as immune modulating genes *TGFβ* and *LAG3*. Taken together, these analyses define a highly diverse MAIT cell transcriptional landscape shaped by chronic activation, with important parallels with conventional CD4^+^ and CD8^+^ T cells.

### Acute MAIT cell activation induces a cytotoxic phenotype marked by GZMB, IFNG, and TNF

We next asked how the prevalence of these populations varied across stimuli and donors during acute and chronic activation (Fig. 2d-e and Supplementary Fig. 2b). In particular, C1 “MAIT1 cytotoxic effector”—which expressed high levels of *IFNG*, *GZMB*, and *TNF*—was enriched by TCR stimulation conditions with 5-OP-RU or αCD3/CD28 compared to cells incubated with IL2 alone (Fig. 3a, b). To validate and further explore these early transcriptional phenotypes, we performed flow cytometric characterization of MAIT cells under the same acute activation conditions.

**Figure 3.**
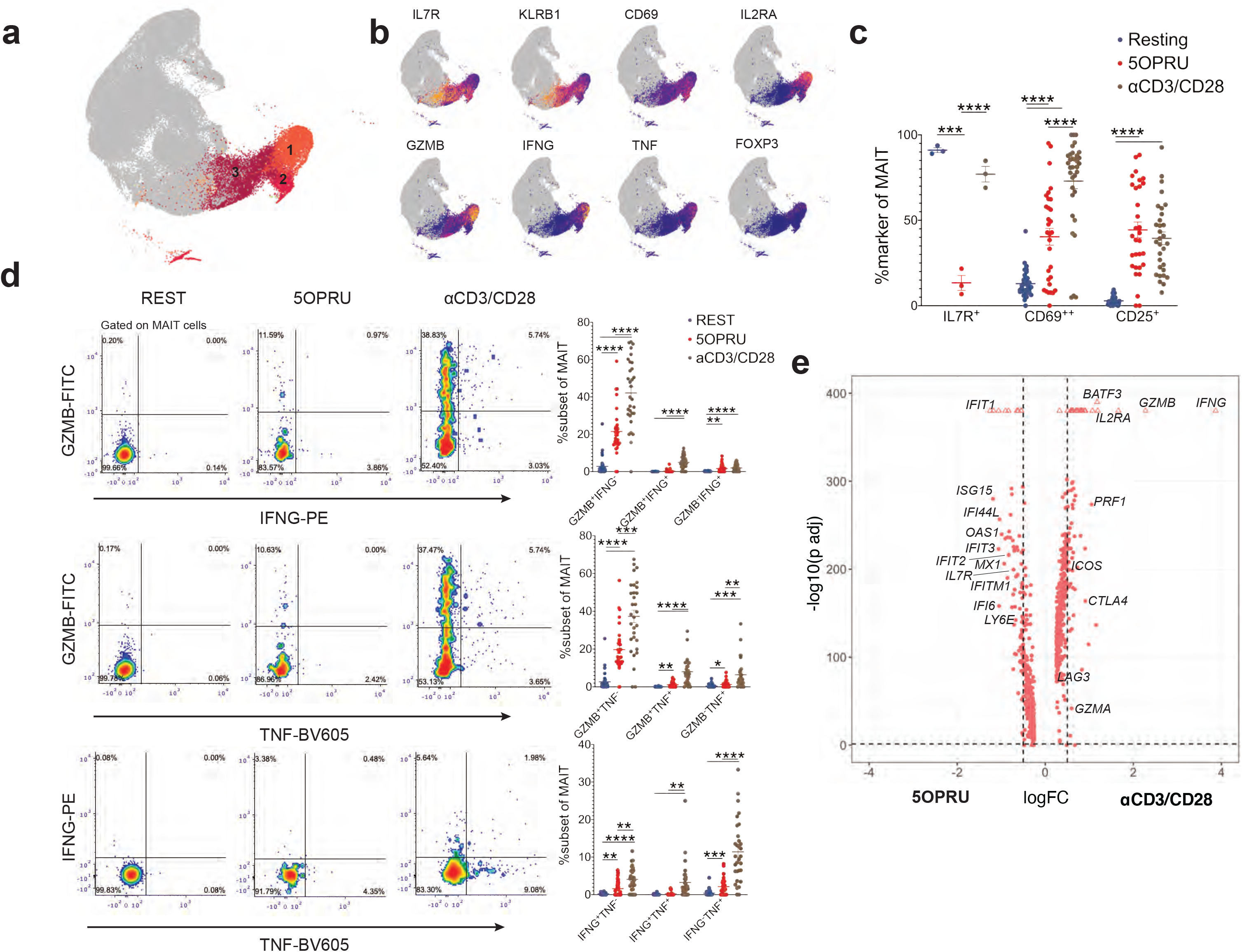
Acute MAIT cell activation induces a cytotoxic phenotype marked by GZMB, IFNG, and TNF. **(a)** UMAP plot highlighting clusters 1-3 enriched in day 1 (D1) incubation conditions. **(b)** UMAP plots of selected MAIT cell marker expression levels at D1 (blue=low expression to yellow=high expression). **(c)** Mean% +/− SEM of MAIT cell extracellular staining of activation markers detected by flow cytometry after 15 hrs of incubation in three conditions: resting (blue), 5OPRU (red), αCD3/CD28 (brown). Results are representative of 4 independent experiments: n=3-30 donors. **(d)** Representative flow cytometry density plots in one donor demonstrating GZMB, TNFα and IFNγ co-staining (left) alongside cumulative plots (right) of mean% +/− SEM for each co-staining condition in multiple donors. Results are representative of 4 independent experiments. n=30 donors. **(e)** Differentially expressed genes in the αCD3/CD28 (+ log fold change; log FC) or 5OPRU (−log FC) conditions at D1. y axis: log FDR-adjusted p value at significance level of FDR P < 0.05. Flow cytometric statistical comparisons were made by unpaired t-test. *p<0.05 **p<0.005 *** p<0.0005 ****p<0.0001; GZMB: granzyme B

First, we confirmed the differential expression of common T cell surface activation markers IL7R, CD69, and CD25 (Fig. 3c). Unexpectedly, IL7R was significantly downregulated by 5-OP-RU, but not αCD3/CD28. CD69 and CD25 were significantly upregulated by both stimulation conditions, though CD69 was more highly expressed with aCD3/CD28, likely indicating the relative potency of this non-specific T cell stimulus. These results demonstrate that MAIT cells rapidly adopt distinct immune phenotypes after acute antigen-specific and non-specific TCR stimulation.

As *GZMB*, *IFNG*, and *TNF* were significantly upregulated in early activation clusters C1 and C2, we next asked if these markers were co-expressed at the protein level. A representative gating strategy of intracellular co-staining for GZMB, IFNγ and TNF is presented alongside cumulative data from 30 healthy donors (Fig. 3d). These analyses confirmed the presence of a MAIT cell population with a combined granzyme and Th1-associated cytokine phenotype enriched with acute activation, consistent with “MAIT1 cytotoxic” C1 enriched in the αCD3/CD28 condition in the single cell RNA-seq analysis. The primary acute activation response in MAIT cells was upregulation of granzyme B alone, which was increased in αCD3/CD28 relative to 5-OP-RU conditions (Fig. 3d). 5-OP-RU and αCD3/CD28 both induced significant co-expression of GZMB and TNFα, though increased co-expression was observed with αCD3/CD28 (Fig. 3d). Only αCD3/CD28 induced IFNγ and TNFα co-expression (Fig. 3d). These results are consistent with increased expression of IFNγ and TNFα with non-specific TCR stimulation.

We next performed a differential expression analysis between MAIT cells stimulated with 5-OP-RU and aCD3/CD28 at D1 to define the gene expression programs associated with antigen-specific or non-specific TCR stimulation (Fig. 3e and Supplementary Table 4). Although both acute TCR-stimulation conditions induced similar transcriptional states (Fig. 2d), we did detect stimulus-specific gene expression patterns, with αCD3/CD28 robustly upregulating *GZMB*, *PRF1*, *IFNG*, *IL2RA*, *BATF3*, *CTLA4*, and *LAG3*. In contrast, 5-OP-RU induced a strong type 1 interferon signaling signature with upregulation of *IFIT1*, *ISG15*, *IFI44L*, *IFIT3*, *IFITM1* and *IFI6*. These findings are consistent with distinct MAIT cell transcriptional programs driven by acute antigen-specific signals versus non-specific T cell stimulation.

### MAIT cells upregulate FOXP3 during acute and chronic activation

Given differential expression of *FOXP3* observed in C2, we sought to further characterize peripheral FOXP3 expressing MAIT cell subsets by flow cytometry. Although few FOXP3^+^ MAIT cells were detected in resting cells, FOXP3 protein was significantly upregulated with 5-OP-RU and αCD3/CD28 during acute stimulation (Fig. 4a,b). In contrast, only 5-OP-RU significantly upregulated FOXP3 during chronic stimulation (Fig. 4b).

**Figure 4.**
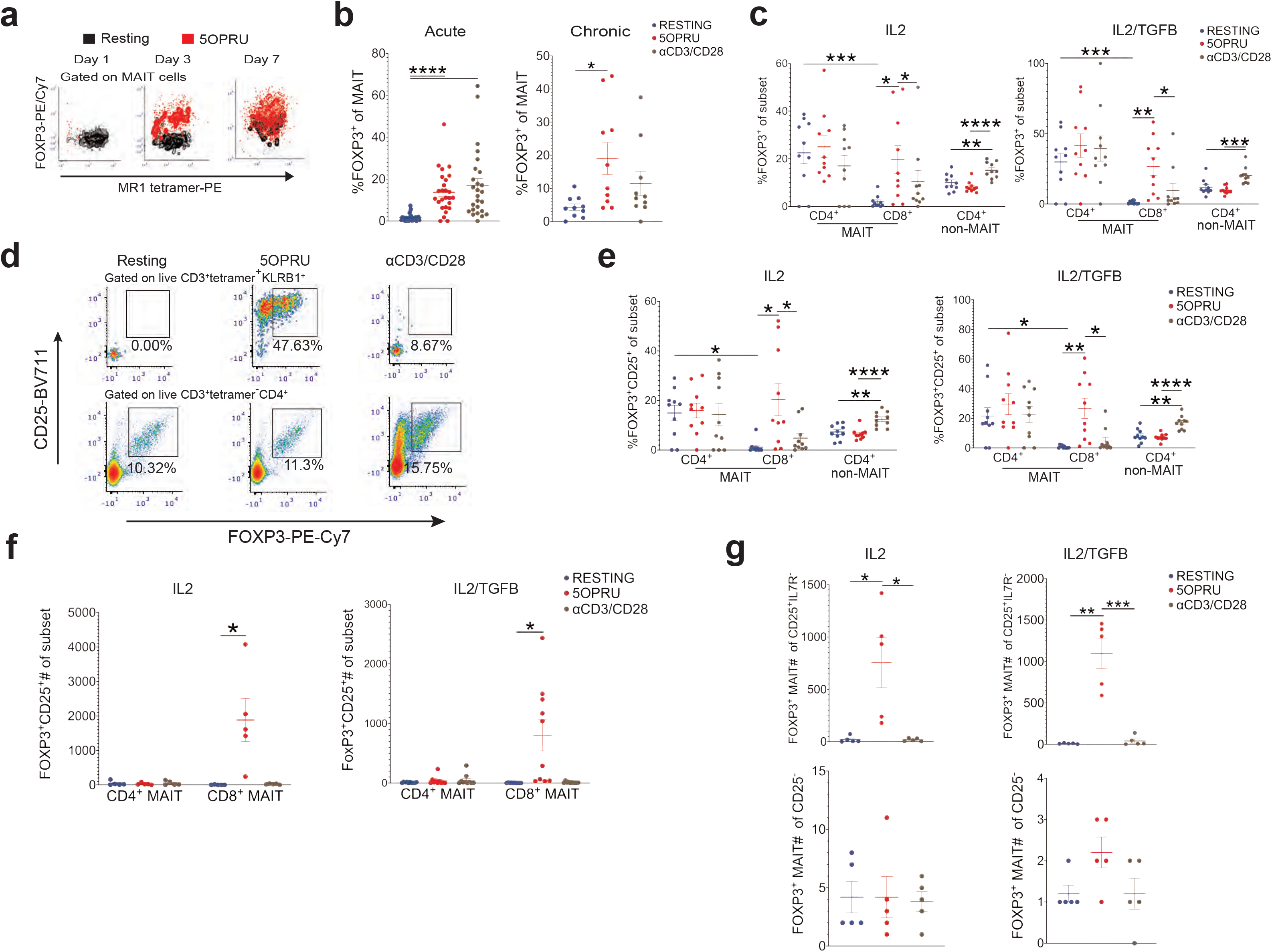
MAIT cells upregulate FoxP3 during acute and chronic activation. **(a)** Representative flow cytometry contour plots of MAIT cell FOXP3 intracellular staining in resting or 5OPRU conditions after 1, 3, or 7 days of stimulation (black=resting; red=5OPRU). **(b)** Mean% +/− SEM of intracellular MAIT cell FOXP3 staining after 1 (D1; acute) or 7 (D7; chronic) days of incubation under resting (blue), 5OPRU (red), or αCD3/CD28 (brown) conditions. Data representative of 3 independent experiments. n=10-26 donors. **(c)** Mean% +/− SEM of intracellular FOXP3 staining in CD4^+^ and CD8^+^ MAIT cells compared to non-MAIT CD4^+^ T cells after 7 days of incubation in the same conditions as (b). **(d)** Representative flow cytometry density plots from one donor demonstrating FOXP3^+^CD25^+^ MAIT cells (top row) or non-MAIT CD4^+^ T cells (bottom row) after 7 days of incubation in the same conditions as (b). **(e)** Mean% +/− SEM of FOXP3^+^CD25^+^ CD4^+^ or CD8^+^ MAIT cells compared to non-MAIT CD4^+^ T cells after 7 days in the same conditions as (b). Cytokine co-incubation conditions included IL2 (left) or IL2/TGFβ (right). Representative of two independent experiments. n=10 donors. **(f)** Mean absolute number +/− SEM of FOXP3^+^CD25^+^ CD4^+^ or CD8^+^ MAIT cells in the same incubation conditions as (e). Representative of 1 independent experiment. n=5 donors. **(g)** Mean absolute number +/− SEM of FOXP3^+^ MAIT cells among CD25^+^IL7R^−^ MAIT cells or CD25^−^ MAIT cells after 7 days in the same incubation conditions as (e).

As FOXP3 is a transcriptional regulator of CD4^+^ regulatory T (Treg) cells and often co-expressed with CD25(Liston et al., 2008; Yamaguchi et al., 2011), we hypothesized that MAIT cells could share similar staining characteristics. First, we compared FOXP3 expression in CD4^+^ and CD8^+^ MAIT cells with non-MAIT CD4^+^ T cells (Fig. 4c). While CD4^+^ MAIT cells had increased FOXP3 expression at baseline, only chronic 5-OP-RU-activated CD8^+^ MAIT cells stably upregulated FOXP3 (Fig. 4c). The addition of TGFβ did not significantly affect MAIT cell FOXP3 expression, though there was a trend for increased FOXP3^+^ expression with TGFβ co-stimulation (Fig. 4c). We next compared FOXP3/CD25 co-expression in MAIT cells with non-MAIT CD4^+^ T cells and found that 5-OP-RU-activated MAIT cells strongly upregulated FOXP3, but CD3/CD28 stimulation only upregulated FOXP3 in conventional CD4^+^ T cells (Fig. 4d). In contrast to Treg cells, the FOXP3^+^CD25^+^ MAIT cells were strongly induced in the CD8^+^ MAIT cell subset after 5-OP-RU activation (Fig. 4e). As previously reported(Hoffmann et al., 2004), αCD3/CD28 stimulation alone resulted in significant Treg cell expansion (Fig. 4e). Analysis of absolute numbers of FOXP3^+^CD25^+^ MAIT cells demonstrate that FOXP3^+^CD25^+^ MAIT cells were most prevalent within the CD8^+^ MAIT cell subset (Fig. 4f) and that the vast majority of FOXP3 expressing cells were detected among CD25^+^IL7R^−^ cells and rarely among CD25^−^ cells (Fig. 4g).

### Chronic activation of MAIT cells expands homeostatic subpopulations while also inducing proliferative, cytotoxic, regulatory, and exhausted phenotypes

We next examined clusters enriched during chronic activation (Fig. 5a). These clusters were broadly associated with expression of *MKI67*, *GZMA*, *GZMB*, *TNF*, *CTLA4*, *ICOS*, *LAG3*, *FAS*, and *PDCD1* (Fig. 5a-b). Despite largely overlapping transcriptional states observed between chronic 5-OP-RU and αCD3/CD28 conditions (Fig. 2e), 5-OP-RU significantly upregulated genes associated with lymphoid development (*LTB*), self-renewal (*LST1*), maturation (*CD52*), motility and proliferation (*STMN1*, *ACTG1*) and recruitment of immune cells (*CCL3*, *CCL4*) (Supplementary Table 4). In contrast, αCD3/CD28 upregulated genes associated with cytotoxicity (*GZMB*, *GZMK*), cell stress (*HILPDA*), and exhaustion (*CTLA4*, *LAG3*) (Supplementary Table 4). This granzyme/exhaustion signature observed was largely driven by C12 defined by cells from Donor 1 (Supplementary Fig. 2c).

**Figure 5.**
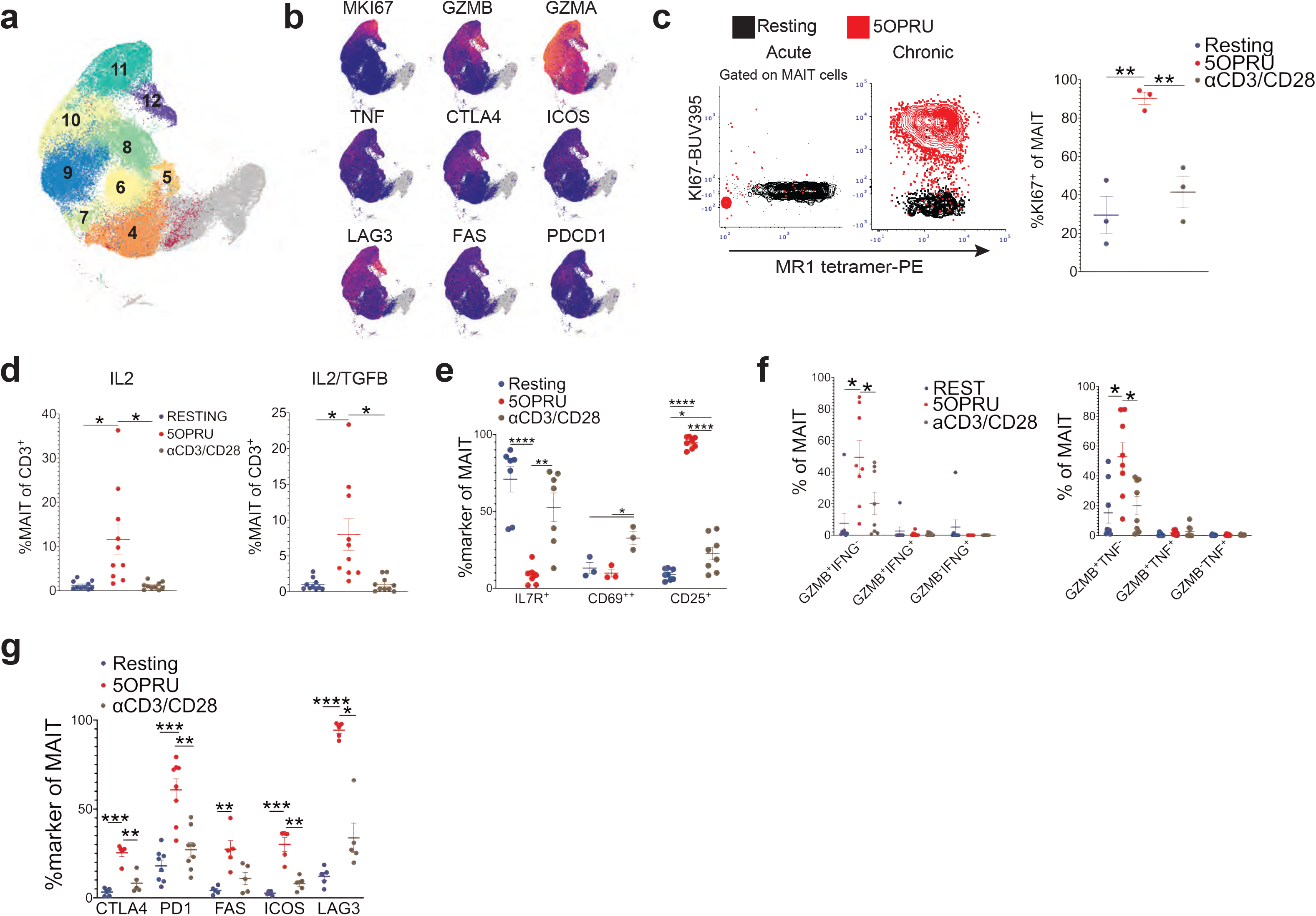
Chronic activation of MAIT cells expands homeostatic subpopulations while also inducing proliferative, cytotoxic, immune regulatory and exhaustion phenotypes. **(a)** UMAP highlighting clusters 4-12 defined by day 7 (D7) incubation conditions. **(b)** UMAP plots of selected MAIT cell marker expression levels at D7 (blue=low expression to yellow= high expression). **(c)** Representative flow cytometry contour plots (left) of intracellular ki67 staining after 15 hours (D1; acute) or 7 days (D7; chronic) of incubation at rest (black) or with 5OPRU (red) accompanied by cumulative data of mean% +/− SEM of MAIT cell ki67 staining after 7 days of incubation under resting (blue), 5OPRU (red) or αCD3/CD28 (brown) conditions (right). Data is representative of one independent experiment. n=3 donors. **(d)** Mean% +/− SEM of MAIT cells among total T cells after 7 days of incubation in the same conditions as (c). Data is representative of two independent experiments. n=10 donors. **(e)** Mean% +/− SEM of MAIT cell surface activation markers after 7 days of incubation in the same conditions as (c). Data is representative of two independent experiments. n=3-8 donors. **(f)** GZMB, IFNγ, and TNFα intracellular co-staining demonstrating mean% +/− SEM of each subset after 7 days of incubation under the same conditions as (c). Data is representative of two independent experiments. n=8 donors. **(g)** Mean% +/− SEM of activation/exhaustion marker staining after 7 days of incubation under the same conditions as (c). Data is representative of one independent experiment. n=5 donors. Data representative of one independent experiment. n=3 donors. Flow cytometric statistical comparisons were made by unpaired t-test. *p<0.05 **p<0.005 *** p<0.0005 ****p<0.0001; GZMB: granzyme B

The transcriptional differences between stimuli were further elucidated at the protein level, where proliferation marker Ki67 was significantly upregulated in the 5-OP-RU but not the αCD3/CD28 condition (Fig. 5c,d). This was consistent with robust antigen-specific MAIT cell expansion not observed with chronic αCD3/CD28 (Fig. 5e). Similar to acute activation, we again detected significant downregulation of IL7R at the protein level only in the 5-OP-RU condition, which was associated with significant upregulation of CD25 not observed with chronic αCD3/CD28 stimulation (Fig. 5f). Similar to acute activation, GZMB was the dominant MAIT cell effector protein (Fig. 5g). Activation/exhaustion markers CTLA4, PD1, FAS, ICOS and LAG3 expression were also enhanced with chronic 5-OP-RU stimulation at the protein level (Fig. 5g).

### MAIT cell subtypes are present during human tuberculosis infection

Finally, we assessed whether the MAIT cell phenotypes observed during acute and chronic activation could also be detected during natural human exposure to *Mycobacterium tuberculosis* (*Mtb*). To this end, we analyzed MAIT cell phenotypes in PBMCs collected from household contacts of active TB patients in Port-au-Prince, Haiti compared to healthy unexposed donors from the same community.

First, we focused our analysis on a subset of markers that characterized CD4^+^ MAIT cells in our single cell analyses. We observed a similar differential expression pattern of CD25, CCR7 and OX40 at the protein level in CD4^+^ MAIT cells relative to other subsets (Fig. 6a-c). Moreover, each of these markers was significantly upregulated in contacts recently exposed to *Mtb,* indicating that the acute activation gene expression pattern characteristic of clusters 1-3 in vitro correspond to in vivo MAIT cell subsets in the context of recent *Mtb* exposure. Consistent with our previous work(Vorkas et al., 2018), these findings indicate that CD4^+^ MAIT cells respond to acute *Mtb* exposure.

**Figure 6.**
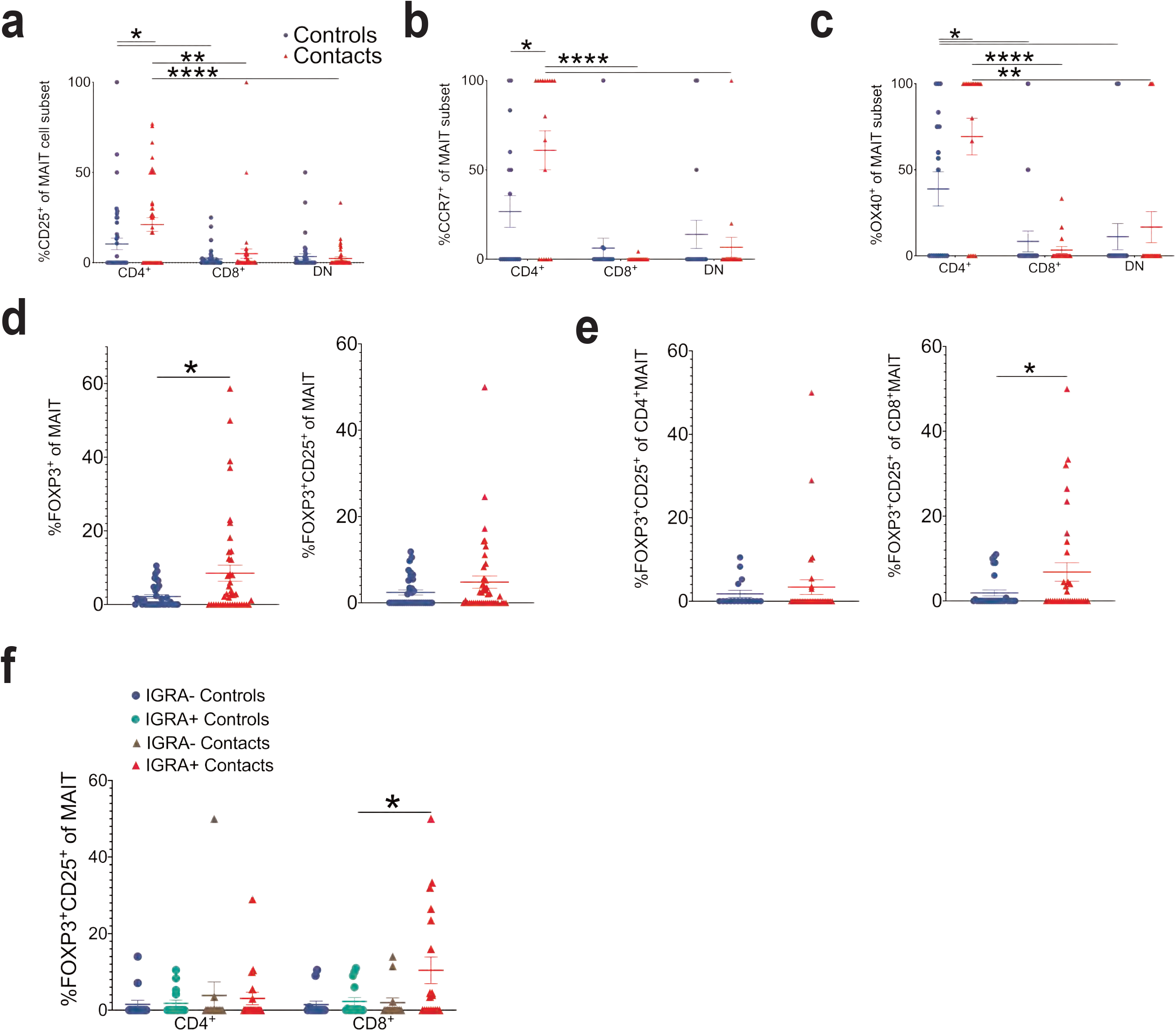
MAIT cell subsets differentially respond to acute and chronic activation by *Mycobacterium tuberculosis*. **(a-c)** Mean% +/− SEM of CD25 (a), CCR7 (b), and OX40 (c) staining of CD4^+^, CD8^+^ or DN MAIT cell subsets in healthy household contacts of TB patients (red triangles) compared to healthy unexposed community controls (blue circles). Same legend also applies to panels (d) and (e). n=18-41 donors. **(d)** Mean% +/− SEM of FOXP3^+^ (left) or FOXP3^+^CD25^+^ (right) MAIT cells. n=36-42 donors. **(e)** Mean% +/− SEM of FOXP3^+^CD25^+^ CD4^+^ (left) or CD8^+^ MAIT cells (right) in the same donors as (d). **(f)** Mean% +/− SEM of FOXP3^+^CD25^+^ CD4^+^ or C8^+^ MAIT cells stratified by IGRA status in the same donors as (d). Flow cytometric statistical comparisons were made by unpaired t-test. *p<0.05 **p<0.005 *** p<0.0005 ****p<0.0001; IGRA: *Mtb*-specific IFNγ release assay.

We next interrogated MAIT cell FOXP3 expression, which was predominant in the CD8^+^ subset after chronic antigenic stimulation (Fig. 4e,f). We found that recently exposed contacts expressed increased FOXP3 relative to healthy controls (Fig. 6d). Consistent with the scRNA-seq results, FOXP3^+^CD25^+^ MAIT cells were most prevalent in the CD8^+^ subset, and FOXP3^+^CD25^+^ CD8^+^ MAIT cells were significantly enriched in contacts (Fig. 6e). When stratifying by *Mtb*-specific IFNγ release assay (IGRA) status, which distinguishes contacts who were exposed and latently infected (IGRA+) from those who remained uninfected (IGRA-), we observed that FOXP3^+^CD25^+^ CD8^+^ MAIT cells were enriched in latently infected contacts (Fig. 6f), further confirming that this MAIT cell subtype is responding to *Mtb* infection. These MAIT cell phenotypes detected in TB contacts closely resemble the CD8^+^ FOXP3^+^CD25^+^ MAIT cells detected with antigen specific stimulation residing in clusters 1-3 (Fig. 3 and 4) and indicate that these cell states defined by single cell transcriptomics are present during acute human *Mtb* exposure and infection.

## Discussion

MAIT cells have been well-characterized as innate-like, rapid responders to microbial antigens with the capacity to mount cytotoxic responses and secrete Th1/Th17-associated cytokines within hours of antigen recognition(Dias et al., 2017; Dusseaux et al., 2011; Vorkas et al., 2018). However, less is known about the transcriptional programs that define MAIT cell adaptation during acute and chronic activation states. Here we provide the first single cell transcriptional atlas of over 76,000 human MAIT cells in homeostatic and acute/chronic activation states coupled with flow cytometric and functional characterization. Our analysis demonstrates distinct transcriptional programs defined by incubation period and quality of stimulus, revealing novel MAIT cell subpopulations not yet described by bulk RNA-seq or traditional flow cytometric approaches.

First, our work contributes further evidence to support divergent transcriptional and functional phenotypes associated with human MAIT cell CD4 or CD8 co-expression. Several prior studies have used CD8 as a prerequisite for defining MAIT cells or did not distinguish between subsets(Coulter et al., 2017; Estevez et al., 2020; Koay et al., 2019; Slichter et al., 2016). Differential expression analysis revealed that CD4^+^ MAIT cells upregulated *IL7R*, *CD25*, *CCR7*, *SELL* along with TNF superfamily receptors *TNFRSF4* and *TNFRSF18*, suggesting that CD4^+^ MAIT cells respond to distinct co-stimulatory signals from the CD8^+^ subset. We hypothesize that this CD4^+^ MAIT cell signature may confer unique tissue-homing functions through *CCR7* and *SELL*. Future studies will help to evaluate whether this CD4^+^ MAIT cell signature confers the capacity to exit the circulation and migrate into tissues.

Recent work in mice and humans shows that CD4 and the transcription factor, LEF1, are co-expressed during MAIT cell intrathymic development with reciprocal downregulation of CD319 (SLAMF7)(Koay et al., 2019). CD4 is subsequently downregulated as MAIT cells mature with reciprocal upregulation of CD8 and CD319 during thymic egress. In a parallel observation in human peripheral blood MAIT cells, we observed that CD4^+^ MAIT cells significantly upregulated *LEF1*, while *SLAMF7* was significantly differentially expressed in CD8^+^ MAIT cells. We hypothesize that the peripheral CD4^+^ MAIT cells may comprise newly generated cells emigrating from the thymus(Fink and Hendricks, 2011). Our findings motivate further study of how MAIT cell activation conditions can select for CD4 or CD8 co-expression in peripheral circulating populations, and further, how these subsets differentially respond to major pathogens. Indeed, our TB data demonstrate that CD25, CCR7 and OX40 are significantly upregulated in CD4^+^ MAIT cells of TB contacts, suggesting that this subset responds to acute *Mtb* exposure.

The second aim of our study was to distinguish between transcriptional phenotypes induced during acute and chronic activation and to characterize functional programs associated with MR1 ligand vs non-specific T cell stimulation. We observed that acutely activated MAIT cells with either stimulus were driven towards similar states defined by a rapid upregulation of GZMB, IFNγ and TNFα. At both the transcript and protein level, this signature was more potently induced by αCD3/CD28, which is possibly explained by non-specific stimulation of all T cells within PBMCs triggering cytokine release by MAIT and non-MAIT T cells alike(Trickett and Kwan, 2003). αCD3/CD28 may mimic other non-specific T cell stimuli derived from whole bacteria or cytokines, which have previously been shown to significantly upregulate GZMB and IFNγ in MAIT cells relative to 5-OP-RU alone(Lamichhane et al., 2019; Suliman et al., 2019). In contrast, acute activation with 5-OP-RU induced a strong type I interferon-driven gene expression phenotype along with T cell maturation marker CD52. Co-stimulation of MAIT cells with 5-OP-RU and type 1 interferons have previously been shown to enhance GZMB and IFNγ release(Lamichhane et al., 2020). Our results demonstrate that acute antigen stimulation alone also enhances MAIT cell type 1 interferon immunity, which may promote survival and proliferation observed during chronic stimulation(Welsh et al., 2012) as well as modulate immunity against bacterial or viral pathogens.(Trinchieri, 2010)

While previous studies have used chronic 5-OP-RU co-stimulation to expand MAIT cells for downstream use during in vitro functional assays(Boulouis et al., 2020; Gherardin et al., 2018a), this is the first investigation of the transcriptional landscape of chronic antigen-specific activation during this proliferative phase. While we observed largely overlapping transcriptional signatures between 5-OP-RU and αCD3/CD28 activation conditions, 5-OP-RU significantly upregulated genes associated with T cell development, self-renewal, proliferation and recruitment of inflammatory cells. In contrast, chronic αCD3/CD28 activation upregulated genes associated with cytotoxicity, stress and exhaustion. These differences between antigen and mitogen stimulation were recapitulated by flow cytometry. Taken together, these results demonstrate that antigen-specific stimulation is required for MAIT cell survival, expansion and sustained effector function during the chronic phase of stimulation.

Moreover, we describe a FOXP3^+^ peripheral blood MAIT cell subset enriched in acute activation cluster C1 with abundant FOXP3 detected at the protein level during chronic 5-OP-RU stimulation. We observed that peripheral FOXP3^+^ induction was detected almost exclusively in the CD8^+^CD25^+^IL7R^−^ subset, in contrast to a recent study(Ling et al., 2016). This surface co-expression of FOXP3 and CD25 in CD8^+^ MAIT cells parallels that of CD4^+^ Treg cells and raises the question of whether FOXP3 expression in MAIT cells may confer a similar immune regulatory function(Bhattacharyya et al., 2018). Importantly, we extend our findings of MAIT cell FOXP3 induction in vitro to immune responses of donors recently exposed to *Mtb*. We found that CD8^+^FOXP3^+^CD25^+^ MAIT cells accumulated after acute *Mtb* exposure in healthy household TB contacts and were significantly more abundant in recently infected donors (IGRA+ contacts). Our results suggest that CD8^+^FOXP3^+^CD25^+^ MAIT cells respond to acute *Mtb* infection.

In addition to FOXP3 expression, our analysis identifies several alternative immune regulatory mechanisms mediated by MAIT cells. We observed that activation/exhaustion marker LAG3 was significantly upregulated in chronically activated MAIT cell clusters (C7, C10, and C12) and was significantly differentially expressed in CD8^+^ and DP subsets. LAG3 can bind peptide-MHC II complexes and inhibit CD4^+^ T cell activation(Maruhashi et al., 2018). Additionally, MAIT cells may target CD4^+^ T cells via MR1 or through secretion of immunomodulatory cytokines like TGFβ. Future studies will help to elucidate the relative contribution of these immune modulation pathways on MAIT cell regulatory activity.

Our study also highlights that MAIT cell activation rapidly drives phenotypic and functional exhaustion. This was evident in both antigen and non-specific stimulation conditions at the transcriptional and protein level. These findings mirror evidence from chronic disease states such as TB, viral hepatitis, and malignancy where increased MAIT cell PD1 expression is associated with peripheral MAIT cell depletion and dysfunction and directly correlates with severity of disease in some studies (Duan et al., 2019; Hengst et al., 2016; Jiang et al., 2020; Leeansyah et al., 2013; Saeidi et al., 2015; Shaler et al., 2017; Yong et al., 2018). While immune checkpoint molecules can be transiently upregulated after T cell activation, we observed sustained expression in chronic activation clusters. At the protein level, these exhaustion markers were significantly upregulated in the 5-OP-RU condition, suggesting that chronic antigen-specific activation strongly induces negative feedback mechanisms to suppress further activation and proliferation. Taken together, these data imply that checkpoint blockade may enhance MAIT cell activity against infectious or malignant disease (Boulouis et al., 2020; Leeansyah et al., 2015; Venken et al., 2018) or contribute to autoimmune complications of immunotherapy(Sasson et al., 2020).

We acknowledge certain limitations to our study. The foremost of these is the small number of individuals in our single cell cohort and donor-to-donor variability. However, the functional flow cytometric validation in multiple donors, including a genetically distinct population from Haiti, strengthens our confidence that the MAIT cell states observed can be applied to other populations. Future studies will test the reproducibility of this transcriptional atlas in additional healthy donors or in specific disease states.

In sum, we present an integrated transcriptional and immune landscape of human MAIT cell activation at single cell resolution. Our results provide a detailed analysis of novel MAIT cell clusters defined by acute and chronic stimulation, including distinct CD4^+^ and CD8^+^ MAIT cell signatures. We also show that these phenotypes are recapitulated in the setting of acute *Mtb* exposure and infection. This functional map of MAIT cell activation biology will help inform future investigations into how MAIT cells specialize in healthy and disease states.

## Acknowledgments

We would like to thank Ojasvi Chaudhury for his assistance in single-cell RNA-seq libraries preparation. We acknowledge the Immune Monitoring, Flow Cytometry, and Integrated Genomics Operation Core Facilities at the Sloan Kettering Institute, Memorial Sloan Kettering Cancer, NY, NY for their expert consultation and services rendered during this project. We also acknowledge the NIH Tetramer Core (Emory University, Atlanta, GA) for providing MR1 tetramers. This work was supported by the Ludwig Center for Cancer Immunotherapy, the Tri-I TBRU, part of the TBRU Network (U19 AI111143), and P30 CA008748. L.M. acknowledges support by The Alan and Sandra Gerry Metastasis and Tumor Ecosystems Center. C.K.V. was supported by NIH NIAID K08 AI132739.

## Author Contributions

C.K.V. and M.S.G. conceived of the study. C.K.V. conducted the experiments. J.A. and K.L. synthesized 5-A-RU. L.M. conducted inDROP scRNA-seq. D.F.W. recruited the Haitian TB contact cohort. C.K. and C.S.L. analyzed single-cell RNA sequencing data. C.K.V., C.K., C.S.L. and M.S.G. analyzed wrote the manuscript with input from all authors.

## Methods

### PBMC collection

Healthy donor PBMCs were obtained with written informed consent at Sloan Kettering Institute, MSKCC, Rockefeller University or ATCC. De-identified blood samples were processed according to the protocols approved by the Institutional Review Board of MSKCC and Rockefeller University. TB household contacts and unexposed community controls were recruited at GHESKIO centers, Port-au-Prince, Haiti (U01AI069421). PBMCs were isolated from peripheral blood using Ficoll-Prep (GE Healthcare) or cell preservation tubes (CPT; Becton Dickinson) according to the manufacturer’s instructions and cryopreserved using 90% fetal bovine serum (Gibco) 10% DMSO (Sigma Aldrich) and stored in liquid N_2_ prior to thawing for in vitro assays.

### PBMC culture

PBMCs were cultured for 15 hrs or 7 days in RPMI (GIBCO) supplemented with 10% (v/v) heat inactivated FBS, penicillin/streptomycin (100 U/mL), L-glutamine (2 mM), sodium pyruvate (1 mM), nonessential amino acids (0.1 mM), HEPES buffer (10 mM) and 2-mercaptoethanol (50 μM) (from GIBCO or Sigma Aldrich) at 37° C 5% CO2 in 96 well plates in the presence of IL2 (GIBCO 100 IU/mL or Roche 1 ng/mL). Additional conditions included 1 ng/mL TGFβ (GIBCO). 2 μM 5-A-RU was added directly to 50 μM methylglyoxal to form 5-OP-RU for in vitro stimulation assays. 5-A-RU was synthesized as previously described(Li et al., 2018), resuspended in 200 μM aliquots, cryopreserved, and thawed as needed. αCD3/CD28 beads (Gibco) were incubated with PBMCs at a 1:2 bead:cell ratio.

### Flow cytometry

Cells were washed once in FACS buffer then incubated with Fc Receptor antibodies (eBioscience; Fisher catalog # 14-1961-73) for 15 minutes at room temperature prior to staining. Extracellular staining was performed at room temperature for 15 minutes in 50 μL FACS buffer (eBioscience) while intracellular staining was performed after 2 hours of incubation in Brefeldin A at 37° C followed by 45 minutes of incubation in 50 μL Permeabilization/Fixation buffer (eBioscience), for 1 hour in 50 μl Permeabilization Buffer (eBioscience) at 4° C

The flow cytometric gating strategy to identify MAIT cells using MR1-5-OP-RU tetramers generated by the NIH tetramer core facility(Corbett et al., 2014; Li et al., 2018) is summarized in Supplementary Fig. 1. The % of MAIT cells of total CD3^+^ cells and the absolute number of cells sorted per condition are presented in Supplementary Tables 1a and b.

Cell staining was performed using the following fluorophore-conjugated antibodies (purchased from BD, eBioscience or Biolegend; clones in parentheses): GZMB (GB11), ki67 (20Raj1), FoxP3 (PHC101), IFNγ (4S.BS3), TNFα (MAb11), CD25 (BC96), OX40 (ACT35), CD161 (DX12), CD3 (UCHT1), CD8 (SK1), CD4 (SK3), EOMES (WD1928), CTLA4 (BNI3), GZMA (CB9), GZMK (GM26E7), IL7R (HIL-7R-M21), CD69 (FN50), ICOS (ISA-3), FAS (DX2), LAG3 (11C3C65) and PD1 (EH12.2H7), CCR7 (3D12).

### Flow-assisted cell sorting and flow cytometric analysis

Stained cells were sorted on a BD Aria sorter or analyzed on a BD Fortessa analyzer. Flow cytometry plots and analysis were performed using FCS Express version 7 (De Novo Software, Pasadena, CA). Graphs and statistical analysis of flow cytometry data were generated using Prism version 11 software (Graphpad Software, San Diego, CA).

### Cell barcoding and scRNA-seq library preparation

The FACS-sorted live-cell suspensions were encapsulated in microfluidic droplets using a Chromium instrument (10X genomics) and reagents provided in Single Cell 3’ Reagent Kit (v3). The barcoded-cDNA libraries were prepared according to the manual (CG00183 Rev B). The viability of cells (stained with 0.2% (w/v) Trypan Blue) prior to loading onto the Chromium instrument was 80-98%. Each sample was encapsulated at a final dilution of ~100 cells/μl. Following the reverse transcription step the emulsion droplets were broken, barcoded-cDNA purified with DynaBeads and amplified by 12-cycles of PCR: 98 °C for 180 s, 12x (98 °C for 15 s, 67 ° C for 20 s, 72 °C for 60 s), and 72 °C for 60 s. The 50 ng of PCR-amplified barcoded-cDNA was fragmented with the reagents provided in the kit, purified with SPRI beads and resulting DNA library was ligated to the sequencing adapter followed by indexing PCR: 98 °C for 45 s; 14x (98 °C for 20 s, 54 °C for 30 s, 72 °C for 20 s), and 72 °C for 60 s. The final DNA library was double-size purified (0.6–0.8X) with SPRI beads and sequenced on Illumina Nova-Seq platform (R1 – 26 cycles, i7 – 8 cycles, R2 – 70 cycles or higher) at depth of 220 million reads per sample.

### Single cell RNA-seq analysis

Pre-processing of scRNA-seq fastq files was conducted using cellranger v3.0.2 (10x genomics). scRNA-seq reads were aligned to the hg19 reference genome (ref-version 3.0.0), and the count matrix of cell barcodes by genes used for downstream analysis was generated using the cellranger count function with parameter ‒expect-cells=3000. The raw count matrix for each sample was obtained from the cellranger count filter_matrix output. Briefly, for each sample, cellranger plots the total UMI count against the cell barcode rank in decreasing order of total counts, and filters cell barcodes out of the resulting count matrix based on the inflection point of the plot. This step minimizes the number of empty droplets that are included in downstream analyses. We created an initial count matrix combining all samples from all patients. Following count matrix generation, cells with > 20% of transcripts derived from mitochondrial genes were considered apoptotic, and were thus excluded. Following this step, all mitochondrial genes were filtered out of the count matrix. Ribosomal genes and the noncoding RNAs NEAT1 and MALAT1 were excluded due to prior reports of strong influence on downstream clustering. Genes with mean raw count < 3.0 were removed from the analysis, and we restricted to KLRB1+ cells, resulting in a final count matrix of 76845 cells and xxx genes for downstream analysis. We used Seurat v2.3.4 to perform standard library size and log normalization. The mean library size was 6680 transcripts per cell.

To examine potential batch effects in the data, we computed the prevalence of each cluster out of all cells in each sample, followed by a centered log-ratio (CLR) transformation for compositional data and PCA. This analysis demonstrated that samples clustered by timepoint, not batch (i.e. donor). Differential expression analyses were conducted using the FindMarkers() function in Seurat using a Wilcoxon test. Specifically, we compared each cluster to all other clusters (one vs all), and genes were considered signature genes for a cluster if the log fold change was greater than 0 and FDR P <= 0.05.

---

**Supplementary Figure 1.**
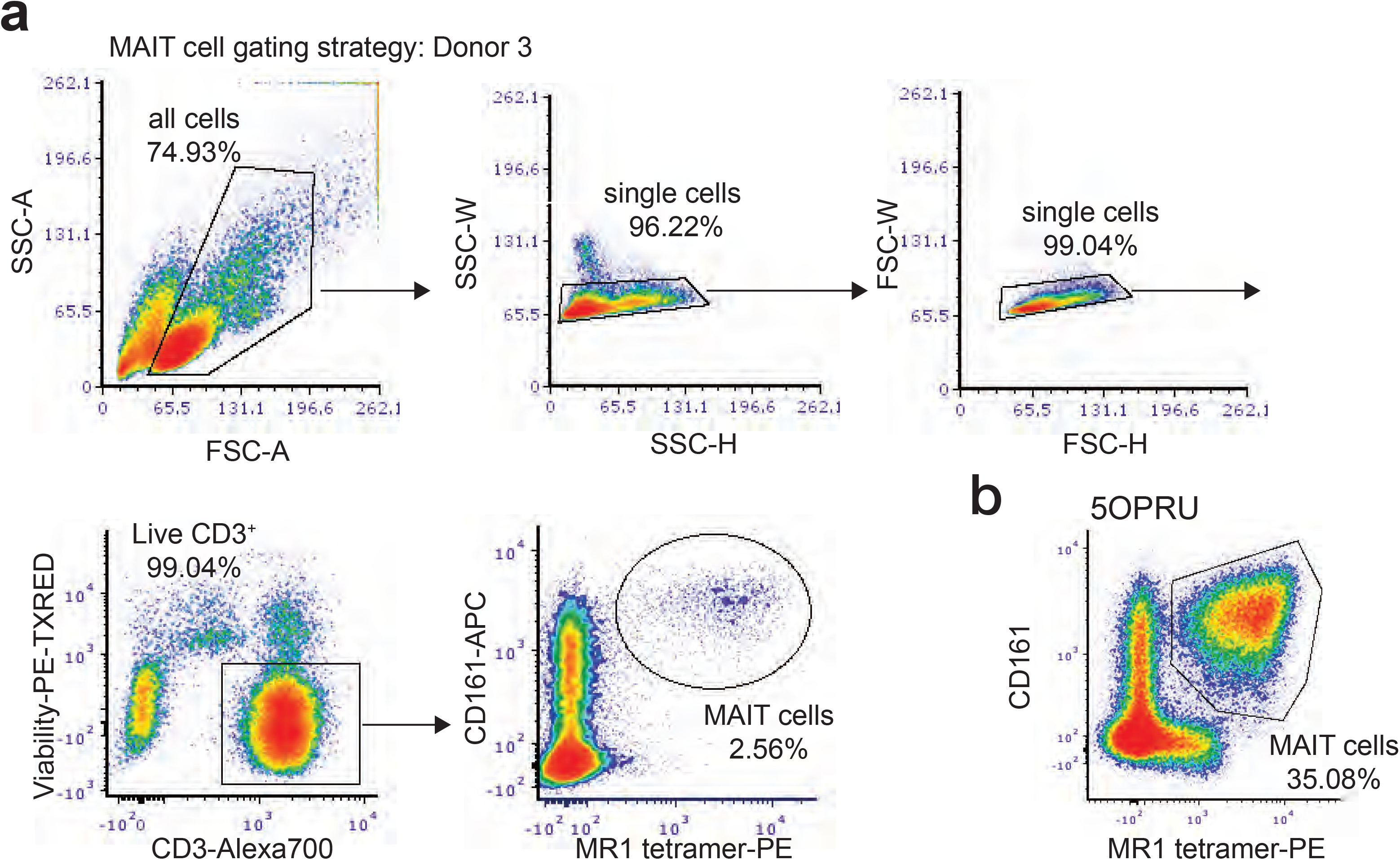
Flow cytometric gating strategy for MAIT cell identification. **(a)** Flow cytometric gating strategy to identify MAIT cells as live, CD3^+^, MR1-5OPRU tetramer^+^, CD161^++^ cells after 15 hrs of overnight rest from thaw in one donor. **(b)** MAIT cell expansion within the CD3^+^ T cell subset after 7 days of 5OPRU incubation.

**Supplementary Table 1.**
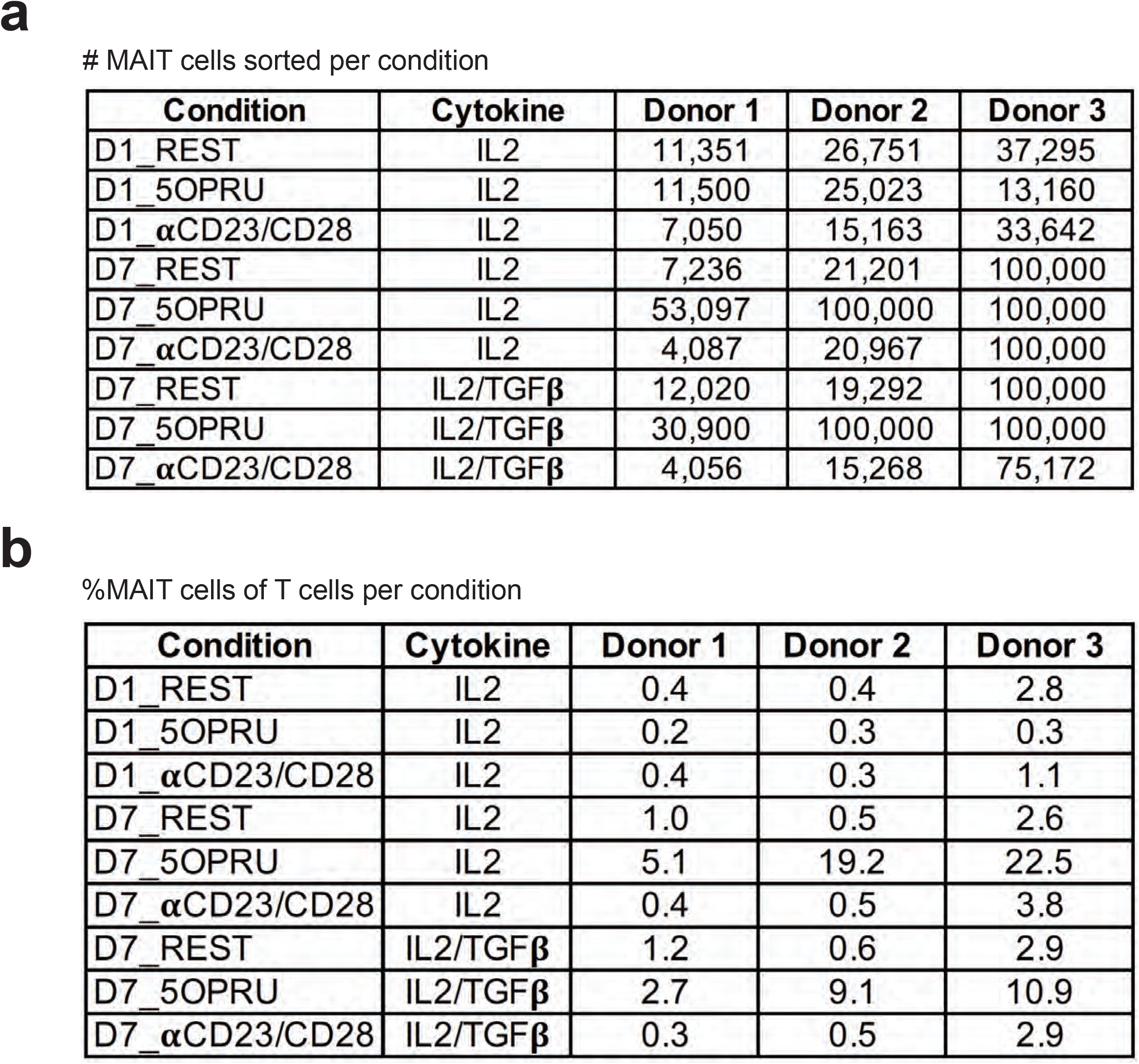
Table of MAIT cell absolute numbers and % of CD3^+^ T cells sorted. Sorted MAIT cells per condition expressed as **(a)** absolute number and **(b)**% of total CD3^+^ T cells.

MAIT cells sorted per condition
| Condition | Cytokine | Donor 1 | Donor 2 | Donor 3 |
| --- | --- | --- | --- | --- |
| D1_REST | IL2 | 11,351 | 26,751 | 37,295 |
| D1_5OPRU | IL2 | 11,500 | 25,023 | 13,160 |
| D1_αCD23/CD28 | IL2 | 7,050 | 15,163 | 33,642 |
| D7_REST | IL2 | 7,236 | 21,201 | 100,000 |
| D7_5OPRU | IL2 | 53,097 | 100,000 | 100,000 |
| D7_αCD23/CD28 | IL2 | 4,087 | 20,967 | 100,000 |
| D7_REST | IL2/TGFβ | 12,020 | 19,292 | 100,000 |
| D7_5OPRU | IL2/TGFβ | 30,900 | 100,000 | 100,000 |
| D7_αCD23/CD28 | IL2/TGFβ | 4,056 | 15,268 | 75,172 |

**Supplementary Table 1. Table of MAIT cell absolute numbers and % of CD3<sup>+</sup> T cells sorted.**
| Condition | Cytokine | Donor 1 | Donor 2 | Donor 3 |
| --- | --- | --- | --- | --- |
| D1_REST | IL2 | 0.4 | 0.4 | 2.8 |
| D1_5OPRU | IL2 | 0.2 | 0.3 | 0.3 |
| D1_αCD23/CD28 | IL2 | 0.4 | 0.3 | 1.1 |
| D7_REST | IL2 | 1.0 | 0.5 | 2.6 |
| D7_5OPRU | IL2 | 5.1 | 19.2 | 22.5 |
| D7_αCD23/CD28 | IL2 | 0.4 | 0.5 | 3.8 |
| D7_REST | IL2/TGFβ | 1.2 | 0.6 | 2.9 |
| D7_5OPRU | IL2/TGFβ | 2.7 | 9.1 | 10.9 |
| D7_αCD23/CD28 | IL2/TGFβ | 0.3 | 0.5 | 2.9 |

**Supplementary Figure 2.**
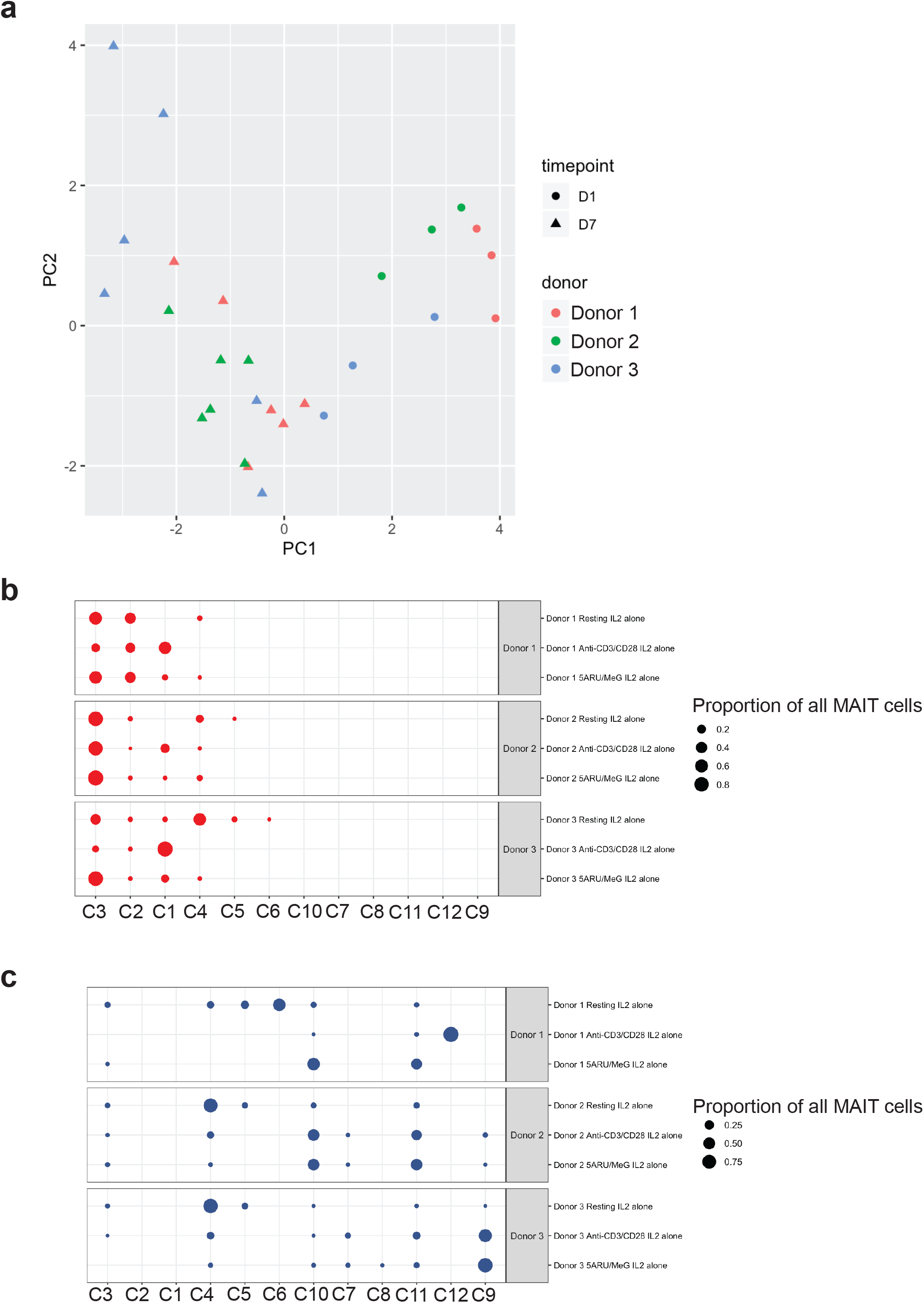
Single cell cluster prevalence per donor and timepoint. **(a)** PCA of CLR-transformed cluster prevalence. Data show broad clustering by timepoint. **(b)** Cluster prevalence per donor and stimulation.

**Supplementary Table 2. Differential expression gene analysis of CD4^+^ versus CD8^+^ MAIT cells in the resting condition at D1 and D7, with all donors pooled together and for individual donors**

**Supplementary Table 3. Differentially expressed genes for each cluster presented in Figure 2.**

**Supplementary Table 4. Differential expression gene analysis of MAIT cells stimulated with 5OPRU or aCD3/CD28 at D1 and D7, with all donors pooled together and for individual donors.**

